# Overcoming pride via the dorsal ACC underlies acceptance of unfair offers

**DOI:** 10.1101/2025.01.09.632093

**Authors:** Shotaro Numano, Chris Frith, Masahiko Haruno

## Abstract

Bargaining is a fundamental social behavior in which individuals often accept unfair offers. Traditional behavioral models, based solely on choice data, typically interpret this acceptance as simple reward-maximization. However, the suppression of emotions such as inequity aversion or pride may also play a critical role in this decision. Incorporating response time alongside choice data provides a means to quantify participants’ internal conflict in suppressing these emotions and deciding to accept unfair offers. In this study, we conducted functional magnetic resonance imaging (fMRI) of the ultimatum game, where participants decided within 10 seconds whether to accept or reject monetary distribution offers from a proposer. Using the drift diffusion model (DDM), we quantified decision-making dynamics based on both choice and response time. Participants who suppressed disadvantageous inequity (DI)-driven rejection (reflected by a lower DDM weight for DI) exhibited heightened dorsal anterior cingulate cortex (dACC) activity in response to DI. Functional connectivity analysis revealed a negative correlation between the dACC and the ventrolateral prefrontal cortex (vlPFC) when DI was large, which encoded both the rejection rates, and the response times associated with accepting DI offers. Furthermore, vlPFC activity was significantly correlated with amygdala activity during high DI conditions, specifically encoding response time for accepting DI offers but not rejection rates. Importantly, these findings could not be captured using standard value-based models that rely solely on choice data. Our results underscore the dACC’s critical role in mediating the suppression of emotional responses to DI, enabling the acceptance of unfair offers in a dynamic bargaining process.

## Introduction

Bargaining is a crucial aspect of human interaction and identifying its cognitive and neural mechanisms is a major objective in human biology. In previous research, a key question for bargaining has been why people reject unfair offers even though accepting any offer is advantageous in terms of their payoff [1,2]. More specifically, several studies reported that the insular cortex is responsive to inequity and contributes to the rejection of unfair offers [3–6]. Other studies have shown that subcortical structures such as the amygdala and ventral striatum also play a key role in representing inequity [7–9] and choosing rejection [8,10–12]. However, much less attention has been paid to the acceptance of unfair offers, which has been regarded mainly as simple reward-maximization [13].

Traditional behavioral studies based on choice (i.e., accepting or rejecting) alone, tend to favor reward-maximization models [13] because these models can succinctly explain observed behavior [13,14]. However, it is also possible that the suppression of emotions plays a central role in accepting unfair offers. Negative emotions could arise from aversion to iniquity. In particular, an unfair, disadvantageous offer could be experienced as disrespectful and an offence to one’s pride.

Consideration of response time alongside choice data may provide valuable information, reveal participants’ internal conflict in suppressing aversion to inequity and choosing to accept unfair offers. Several previous studies have categorized response times as fast or slow [15–18], and analyzed them in the context of dual-process theory [19]. Fast responses are often linked to intuitive decision-making, whereas slow responses are associated with more deliberate thought processes. However, there are limitations in such interpretations. It is difficult to determine whether slow responses stem from a type (i.e., slow or fast) of decision-making process or from conflicts between competing options [18]. Response time may also be influenced by other variables, such as the difficulty in distinguishing between options [15]. It is likely to be more constructive to consider response time and choice simultaneously when examining participant’s dynamic valuation process.

Sequential sampling models, particularly drift diffusion models (DDMs; **Fig 2A**) [20], are well-established computational models that estimate a participant’s binary decision-making process by using both choice and response time. These models posit that individuals accumulate information incrementally during a trial and make a binary decision once enough evidence is gathered. The DDM has four parameters: decision boundary, relative starting point, non-decision time, and drift rate. The decision boundary represents the threshold of information required to make a choice. The relative starting point reflects the participant’s prior response bias. Non-decision time refers to the duration spent on processes unrelated to decision-making, such as encoding the stimulus and executing a motor response like pressing a button. The drift rate reflects the participant’s inclination toward one option (A or B) over time. Although originally introduced in behavioral psychology [21], DDMs have been utilized by neuroscientific studies for both animals and humans [22–25], and applied to a variety of tasks, ranging from perceptual decision-making [25–28] to social behavior [29,30].

In this study, we aimed to uncover the neural dynamics for accepting unfair offers. We conducted an fMRI experiment using the ultimatum game. In each trial, participants decide to accept or reject a money-distribution offer from a proposer within 10s. We analyzed behavior and fMRI data using a DDM. We assume that the drift term relates to self-reward (SR), and advantageous and disadvantageous inequity (AI and DI, respectively). We hypothesize that, if any suppression mechanism of DI-related negative feeling underlies the acceptance of unfair offers, we should observe some DI-correlated brain activity which is large in people who exhibited a small DDM (drift) coefficient for DI. In other words, we think that participants who do not reflect DI in their choice (i.e., they accept unfair offers) but need long response times, are suppressing the DI-related negative feelings in their decision making. On the basis of previous studies on cognitive control [31–35] we hypothesize that the anterior cingulate cortex may be involved in this process.

## Results

### Effects of disadvantageous inequity on choices and response times

Healthy participants (n=63) underwent an fMRI scan while playing an ultimatum game [13,14] as a responder (**Fig 1A**). In this game, proposers made a series of offers about money sharing to the responder, who decided whether to accept or reject each offer within 10 seconds. If the responder accepted an offer, the money was distributed as proposed; otherwise, neither side received any money. Each proposal consisted of pairs of rewards for the participant (self-reward; SR) and the proposers (other-reward; OR).

**Figure 1.**
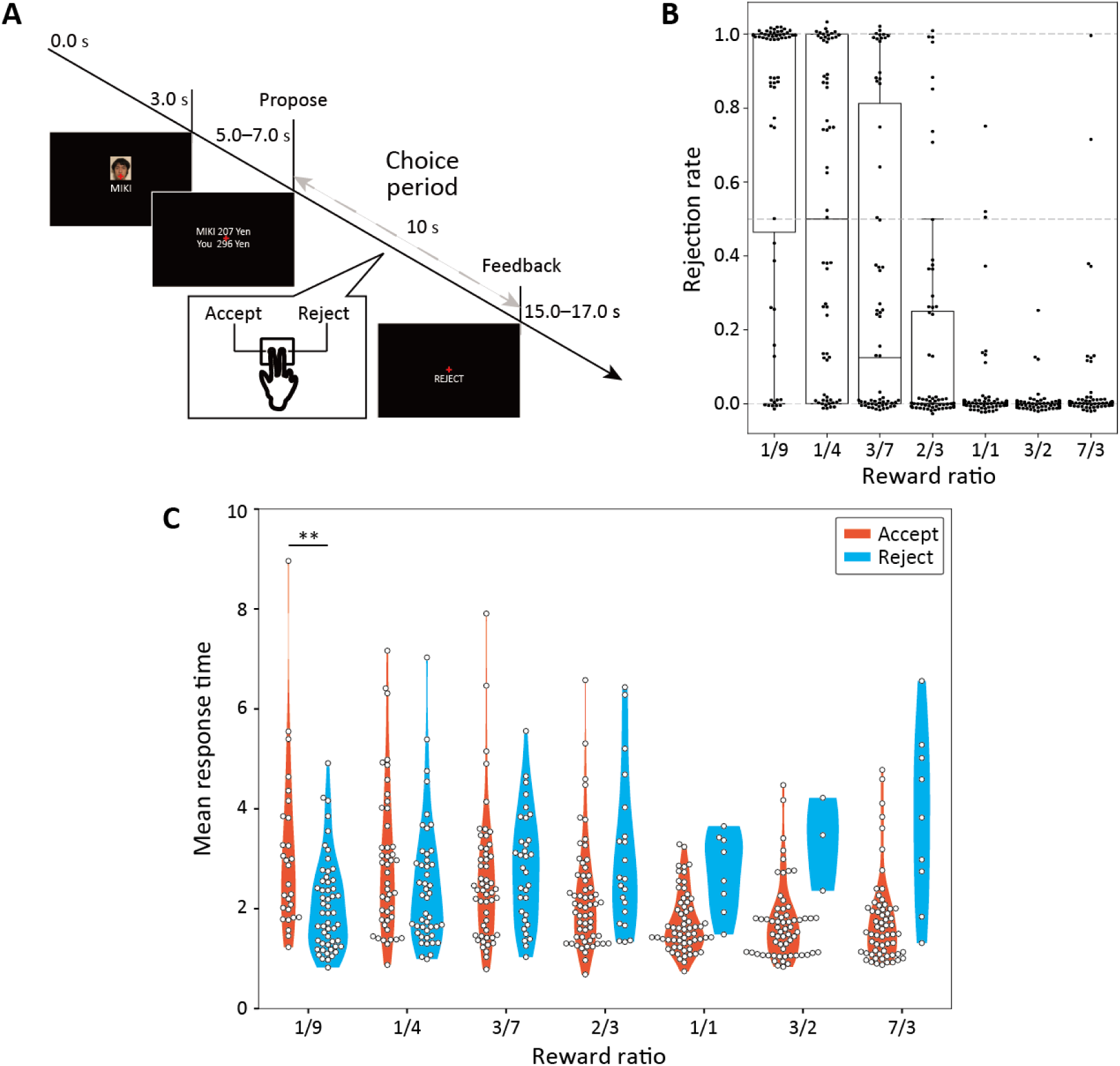
Task and behavioral results (A) Task design. Participants performed the ultimatum game as a responder. In each trial, they chose to accept or reject the displayed proposal of money distribution within 10 s. When the participant accepted the proposal, the money was distributed as proposed. Otherwise, both received no money. (B) Rejection rate for different proposals. Seven different ratios of self- and other- rewards (SR and OR) were used. The rejection rates increased in higher disadvantageous inequity (DI) conditions (e.g., 1/9), and it decreased in lower DI conditions (e.g., 1/1). The rejection rates also increased in higher advantageous inequity (AI) conditions (e.g., 2/3). (C) Response times in different conditions. With the increase in SR, response time for acceptance decreased, while the one for rejection increased. In the highest DI condition (1/9), the mean response time for acceptance was slower than the one for rejection (*t*(36) = 2.97, \*\**p* = 0.00528, Welch’s t-test).

**Figure 2.**
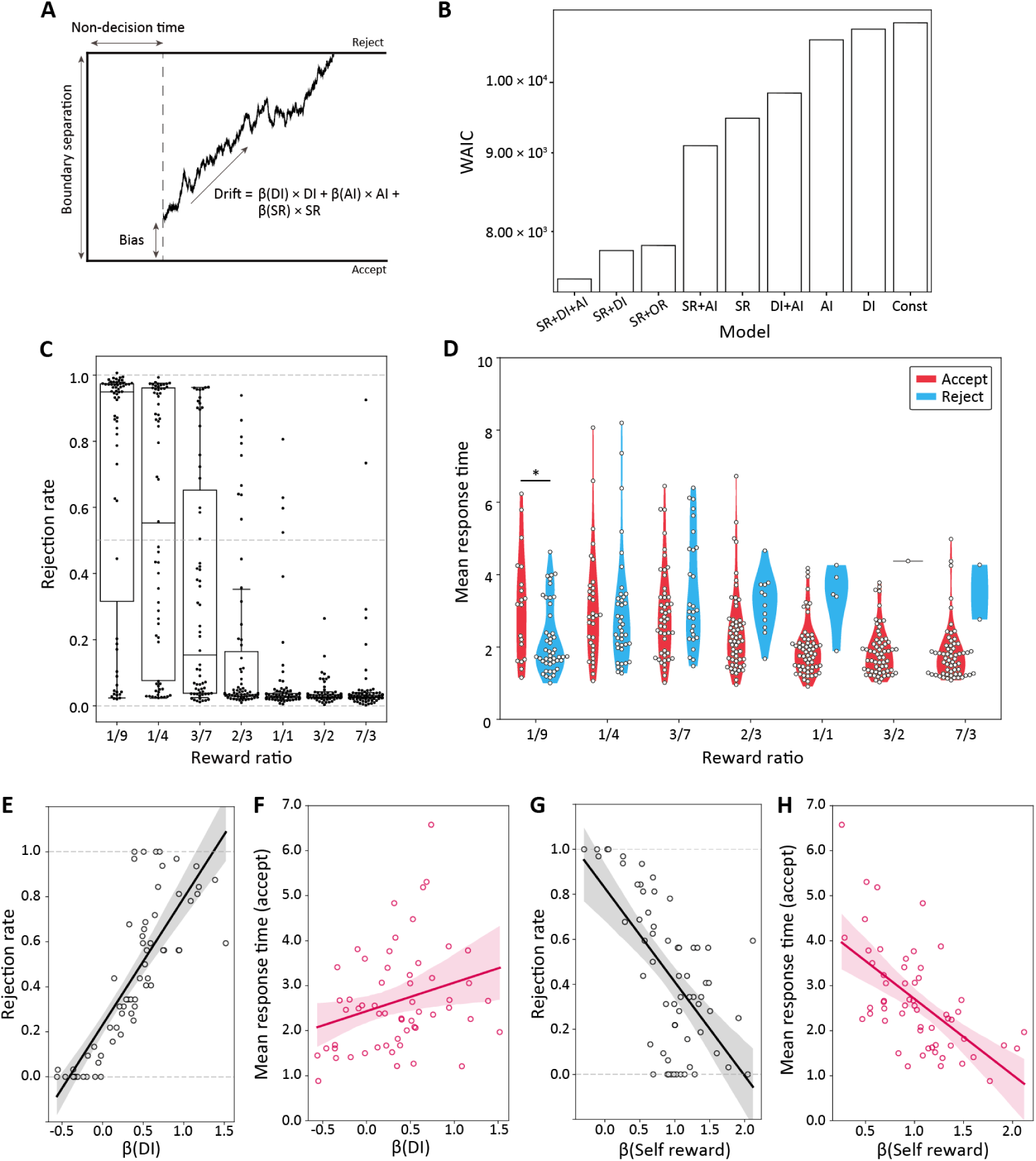
Model selection results and the relationship between drift parameters for disadvantageous inequity and self-reward, and behavior **(A) Drift diffusion model (DDM).** The DDM contains four parameters: drift, non- decision time, boundary and bias. Drift is represented as a liner combination of self- reward (SR), disadvantageous inequity (DI) and advantageous inequity (AI). We selected the best model using the WAIC. **(B) Model selection results.** The best drift term contained SR, DI and AI. Simulated behaviors for (**C) rejection rates, and (D) mean response times for acceptance and rejection in different conditions.** The best model replicated key features in the behavioral data (see also **Fig 1B and 1C**; \**p* = 0.0165 in **D** in the 1/9 condition). **(E)** *β*(**DI**) **and the rejection rate in the DI condition.** The rejection rate increased significantly with increasing *β*(DI) (Pearson’s *r* = 0.819, *p* = 2.38 × 10^−16^ ). (**F)** *β*(**DI**) **and the mean response time for acceptance in the DI condition.** The mean response time increased significantly with increasing *β*(DI) (Pearson’s *r* = 0.274, *p* = 0.0390). (**G)** *β*(**SR**) **and the rejection rate.** The rejection rate increased significantly as *β*(SR) increased (Pearson’s *r* = −0.634 , *p* = 2.41 × 10^−8^). **(H)** *β*(SR) and the mean response time for acceptance (Pearson’s *r* = −0.605, *p* = 6.27 × 10^−7^)

Human behavior in the ultimatum-game has been reported to be influenced by disadvantageous inequity (DI) and advantageous inequity (AI) separately (both defined for SR and OR; see Computational models for inequity-aversion in the Methods) [2,13,36]. We therefore first confirmed that participant behaviors were consistent with prior findings regarding DI and then examined the relationship between the choice and response time (**Fig 1B**). The mean rejection rate for a disadvantageous offer (i.e., SR/OR < 1/1) was 44.0% (s.d. 33.1%), while that for an advantageous offer (i.e., SR/OR ≧ 1/1) was 2.81% (s.d. 9.48%). The difference between the two conditions was significant (*t*(72) = 9.49, *p* = 2.58 × 10^−14^, paired t-test). A repeated one-way analysis of variance (ANOVA) of the mean rejection rates yielded a significant main effect of the SR/OR rate (*F*(6,434) = 52.9, *p* < 7.60 × 10^−49^). The mean rejection rates in the strongest (i.e., SR/OR = 1/9) and weakest (i.e., SR/OR = 2/3) DI trials were 73.5% (s.d. 39.1%) and 17.5% (s.d. 30.9%), with the former significantly higher than the latter (*t*(117) = 8.93 and *p* = 7.00 × 10^−15^ , paired t-test). These results clarified that participants rejected more as the degree of DI increased.

We next examined the participants’ response time (**Fig 1C**). The mean response time for disadvantageous offers was 2.56 s (s.d. 1.30 s) and to advantageous offers it was 1.91 s (s.d. 1.05 s). The mean response time for acceptance was significantly longer than the one for rejection in the strong disadvantageous offers (i.e., SR/OR = 1/9 and 1/4; *t*(123) = 3.48 and \*\*\**p* = 6.82 × 10^−4^, Welch’s t-test). By contrast, for advantageous offers (i.e., SR/OR = 3/2 and 7/3), the mean response time was significantly longer for the rejection than for the acceptance (*t*(12) = 4.34 and \*\*\**p* = 0.00105, Welch’s t-test). These results confirmed that only for strong disadvantageous offers were acceptances slower than rejections.

To explore the dynamic neural processes for accepting unfair offers, we constructed drift-diffusion models (**Fig 2A**; see **Materials and methods**) and selected the best one to predict behavior (**Fig 2B**). More specifically, we calculated the widely applicable information criterion (WAIC; [37,38]) which is a generalization of the Akaike information criterion (AIC) in Bayesian modeling. The best model with the lowest WAIC included three components in the drift term: SR+DI+AI (WAIC = 7.45 × 10^3^). We also found that the model which only included a constant in the drift term had poor predictive accuracy and the SR+DI model worked reasonably well, indicating that the DI term significantly improved the predictability. These results in **Fig 2B** also demonstrated that SR and OR are insufficient to account for choices and response times in the ultimatum game.

We then checked whether the obtained posterior distributions for the best model reproduce the choices and response times. A repeated ANOVA of the posterior prediction of choice (predictive rejection rates; **Fig 2C**) detected a significant main effect of the reward ratio ( *F*(6,434) = 54.5, *p* = 5.68 × 10^−50^ ). Similarly, a repeated two-way ANOVA of the posterior prediction of response time (the predictive mean response time for each choice; **Fig 2D**) had significant main effects of reward ratio and choice and an interaction effect (**Table 2**). The mean response time for acceptance was significantly longer than for rejection in the 1/9 and 1/4 condition (*t*(94) = 2.28, *p* = 0.0249, Welch’s t-test), These results suggest that the best DDM model captured all key aspects of the participants’ choices and response times.

**Table 1.** Repeated two-way ANOVA of the mean response time for each choice.

| Variable | Sum of squares | df | F value | P value |
| --- | --- | --- | --- | --- |
| Reward ratio | 94.2 | 6 | 12.5 | $3.59 \times 10^{-13}$ |
| Choice | 18.3 | 1 | 14.5 | $1.55 \times 10^{-4}$ |
| Reward ratio × Choice | 79.0 | 6 | 10.7 | $5.59 \times 10^{-11}$ |
| Residual | $6.68 \times 10^2$ | 531 | | |

**Table 2.** Repeated two-way ANOVA of the mean predictive response time for each choice.

| Variable | Sum of squares | df | F value | P value |
| --- | --- | --- | --- | --- |
| Reward ratio | 84.9 | 6 | 11.5 | $5.11 \times 10^{-12}$ |
| Choice | 12.2 | 1 | 9.90 | $1.76 \times 10^{-3}$ |
| Reward ratio × Choice | 42.6 | 6 | 5.77 | $8.26 \times 10^{-6}$ |
| Residual | $5.70 \times 10^2$ | 464 | | |

We conducted a correlation analysis between individual parameters of the best model and the mean rejection rate and the mean response time of the same individual (**Fig 2E** to **H**). *β*(DI), which represents the DDM coefficient for DI in the drift term, showed the strongest correlation with the mean rejection rate (**Fig 2E**; Pearson’s *r* = 0.819 and *p* = 2.38 × 10^−16^) and the mean response time for acceptance (**Fig 2F**; Pearson’s *r* = 0.274 and *p* = 0.0390 ) in disadvantageous conditions. We also found that *β*(SR) correlated with the mean rejection rate (**Fig 2G**; Pearson’s *r* = −0.634 and *p* = 2.41 × 10^−8^) and the mean response time for acceptance (**Fig 2H**; Pearson’s *r* = −0.605 and *p* = 6.27 × 10^−7^) in unfair (advantageous and disadvantageous) conditions. These results demonstrated that *β*(DI) accurately encodes both behavioral choices and response times in disadvantageous conditions (see also [55] for more detailed analysis of behaviors and DDM parameters).

### Participants with a smaller *β*(DI) showed larger dACC activity correlated with DI

Having established that *β*(DI) accurately represents behavioral choices and response times in disadvantageous conditions, we used this parameter in our fMRI analysis. First, we conducted a parametric fMRI analysis. As we hypothesized that, for disadvantageous offers, accepting behavior is realized by the suppression of the aversion to DI, we sought the brain activity correlated with DI at the individual level and beta values of the group correlated with −*β*(DI). The assumption behind this analysis was that the suppression process should be stronger when DI is large and even stronger in participants who accept more disadvantageous offers (i.e., correlated with −*β*(DI) ). This analysis identified activity in the dACC (**Fig 3A** and **Table 3**; *p* < 0.05 cluster-level FWE corrected). We also examined BOLD time-series at the peak and confirmed that the five participants who had the smallest β(DI)s and the five participants who had largest β(DI)s showed an increase and a decrease in the dACC activity, respectively (Fig. 3B; they were significantly different, Welch’s t-test, Holm-Bonferroni corrected ** *p* = 0.00296 ). These results suggested that the cingulate area plays a crucial role in the top-down control of DI aversion.

**Figure 3.**
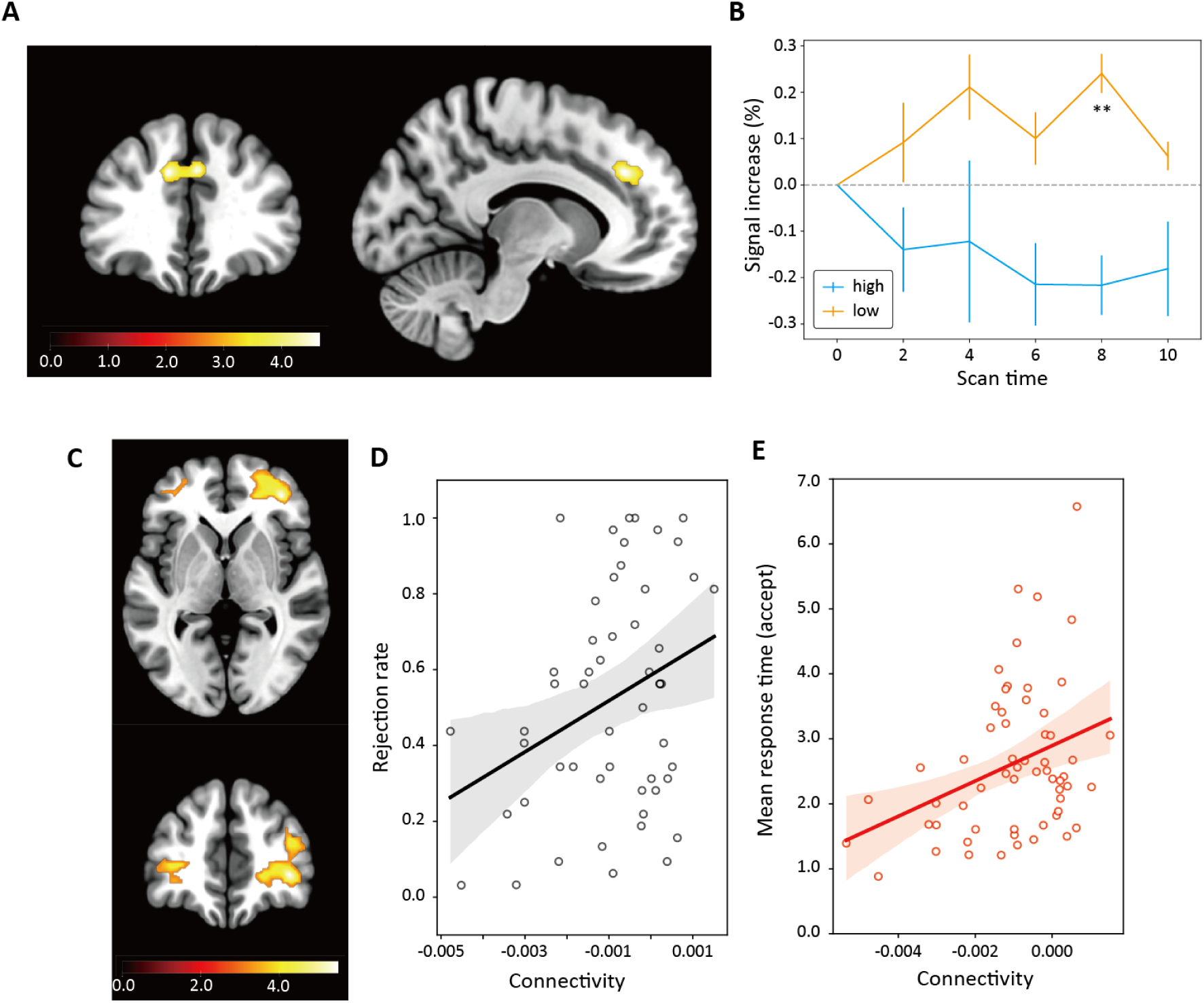
Neural activity correlated with DI in the participants who had a small *β*(DI) and results of the psychophysiological interaction (PPI) analysis. (A) Neural activity correlated with DI weighted by - *β*(DI). We found neural activity in the dorsal ACC (MNI coordinate = [-10, 34, 32]; cluster-level family-wised error (FWE) corrected p < 0.05). We listed other activities in **Table 3**. (B) dACC BOLD signal time-series. The BOLD signal time-series at the peak (MNI coordinate = [-10, 34, 32]) showed an increase and a decrease in the five participants had smallest β(DI)s and the five participants who had largest β(DI)s, respectively (they were significantly different; Welch’s t-test, Holm-Bonferroni corrected ** *p* = 0.00296 ). (C) Functional connectivity between the ACC and vlPFC. The vlPFC (MNI coordinate = [38, 44, 0]; cluster-level family-wised error (FWE) corrected p < 0.05) area had a negative interaction with the ACC when DI was large (see also **Table 4**). (D) Relationship between the connectivity and rejection rates. The mean rejection rate decreased as connectivity became negative (Pearson’s *r* = 0.318, *p* = 0.0201). (E) the connectivity and mean response time for acceptance in the DI condition. The response time for acceptance decreased (became faster) as the connectivity became negative (Pearson’s *r* = −0.340, *p* = 0.00851).

**Table 3.**
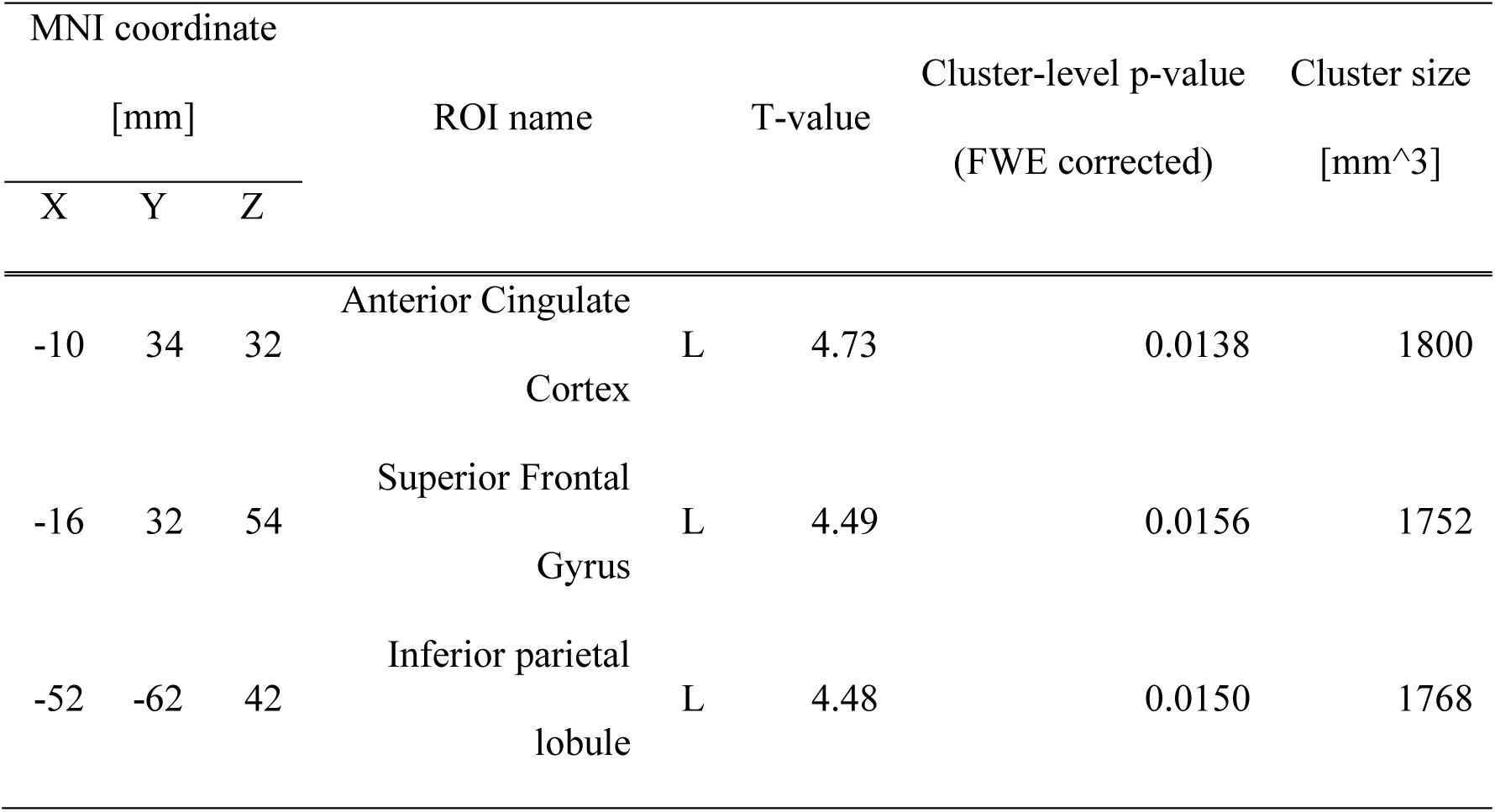
Neural activities to DI weighted by. −*β*(**DI**). We performed cluster-level family-wised error correction (p < 0.05), and all clusters are displayed.

**Table 4.**
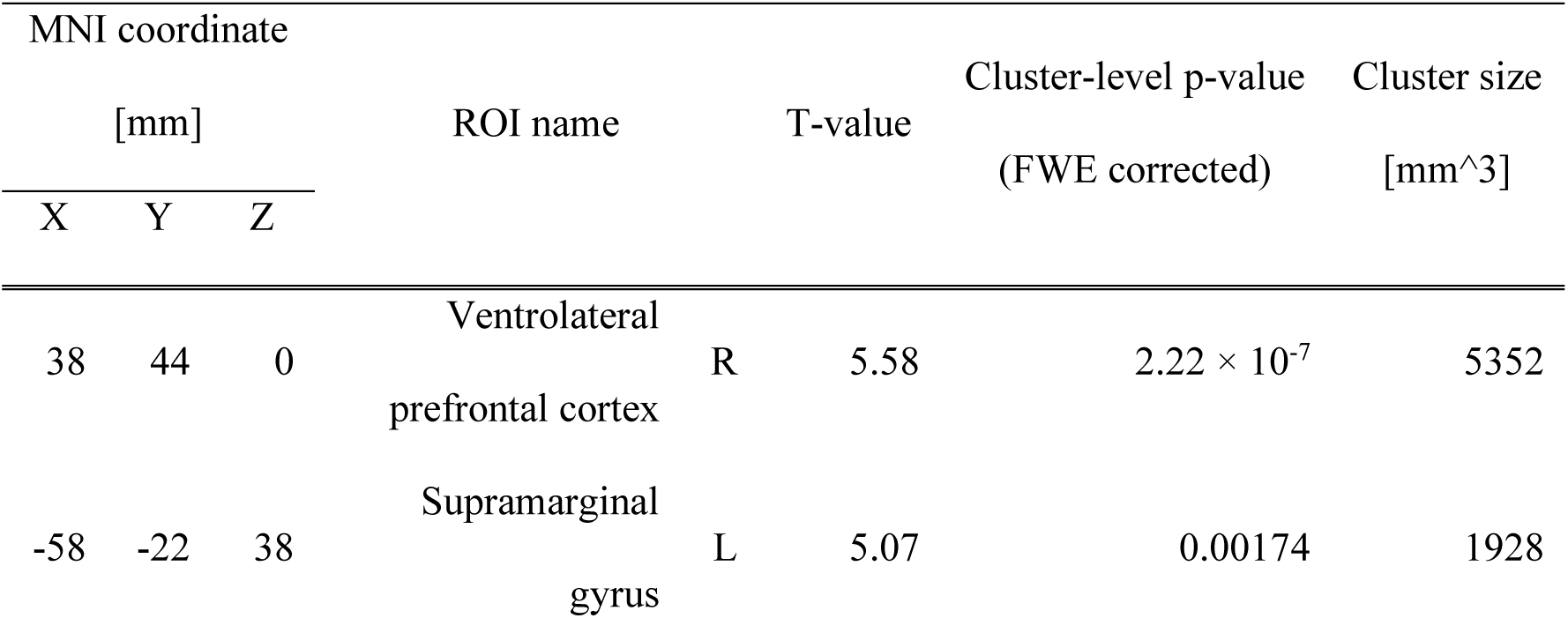
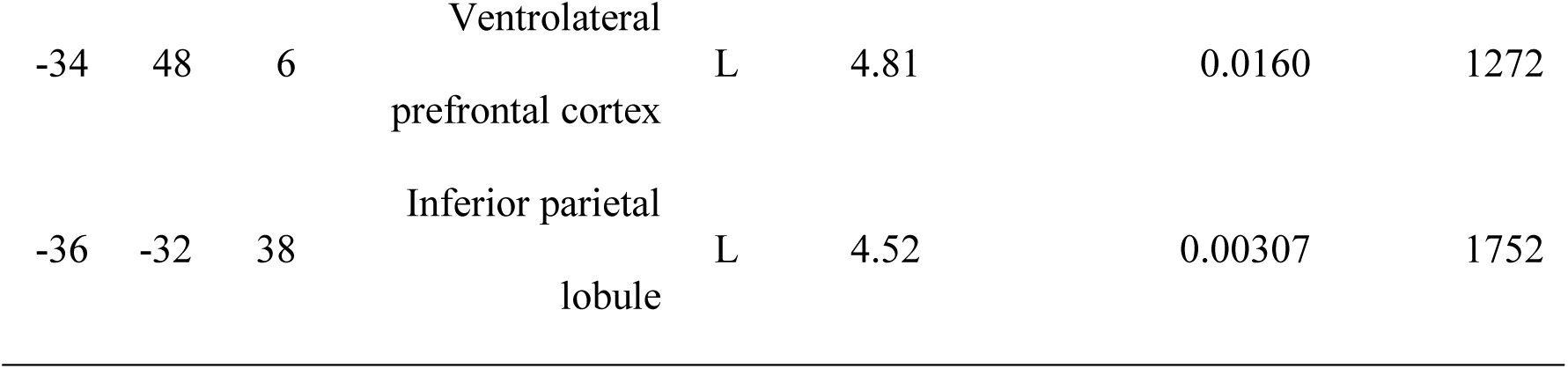
Neural activity identified by the PPI analysis whose seed ROI is the dACC. We performed cluster-level family-wised error correction (*p* < 0.05), and all clusters are displayed.

### dACC-vlPFC interaction encodes rejection rate and response times for accepting disadvantageous offers

To further examine the suppression mechanism by the dACC (**Fig 3A** and **Table 3**), we conducted a psychophysiological interaction (PPI) analysis using the gPPI toolkit [39]. We defined a volume of interest from the peak positions of each cingulate cluster and ran PPI by setting these regions as the seed. We looked for significant negative interactions when DI was large.

We found a significant negative interaction between the dACC (**Fig 3A** and **Table 3**; MNI coordinates = [-10, 34, 32]) and the vlPFC (**Fig 3C** and **Table 4**; cluster-level FWE corrected p < 0.05; MNI coordinates = [38, 44, 0]) when DI was large. Notably, we did not use any DDM-derived parameters to reveal this connectivity because such use of DDM parameters may produce false-positive correlations with response times. This result suggests that the dACC suppresses the vlPFC when DI is large. We did not find any significant connectivity to other seeds in Table 3.

We then tested if this interaction between the dACC and the vlPFC encodes the rejection rate and the response times for accepting disadvantageous offers. We found that the dACC-vlPFC connectivity exhibits a significant correlation with the rejection rate (**Fig 3D**; Pearson’s *r* = 0.318 and *p* = 0.0201 ) and the response time for accepting disadvantageous offers (**Fig 3E**; Pearson’s *r* = −0.340 and *p* = 0.00851). We did not find any significant correlation with rejection rates and response time for other structures in Table 4. These results suggest that suppression of the vlPFC via dACC underlies the decision to accept DI offers.

### vlPFC-amygdala interaction encodes response times for accepting disadvantageous offers

The vlPFC has anatomical and functional connections to subcortical regions such as the amygdala and is involved in emotion regulation [40–42]. Therefore, we hypothesized that the vlPFC has functional connectivity to subcortical regions, particularly amygdala, at every offer timing, and performed a PPI analysis. We defined a volume of interest from the peak positions of vlPFC (MNI coordinates = [38, 44, 0]). The result revealed an interaction between the vlPFC (**Fig 3C**) and the amygdala (**Fig 4A** and **Table 5**; cluster- level FWE corrected p < 0.05 with small volume correction; MNI coordinates = [24, -8, -12]). This result confirmed that the vlPFC and the amygdala are synchronously active at offer timing when DI was large.

**Figure 4.**
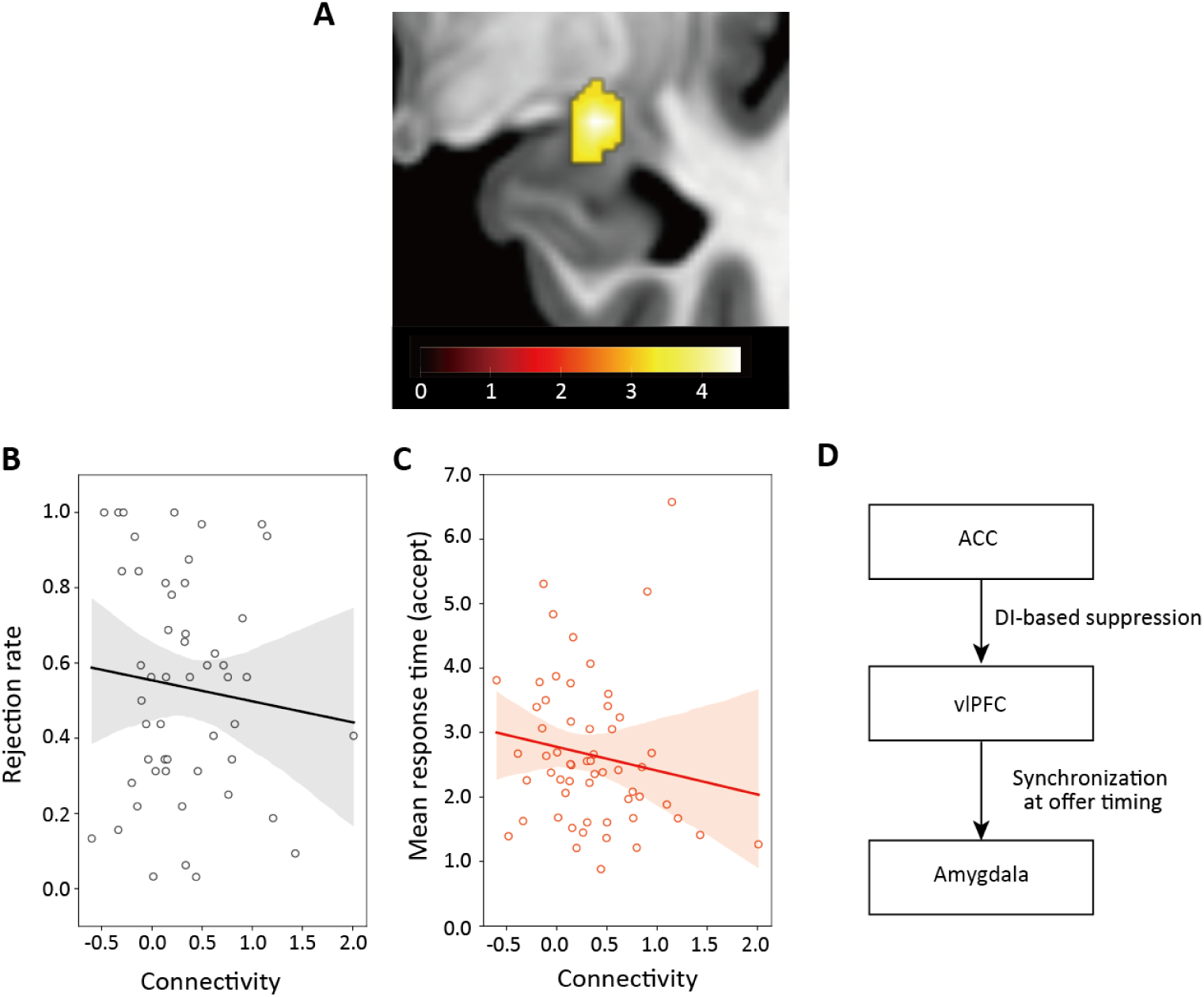
Psychophysiological interaction analysis of the vlPFC (A) Functional connectivity of the vlPFC cluster. The amygdala (MNI coordinate = [24, -8, 12]; uncorrected p < 0.001) had a positive interaction with the vlPFC cluster when DI was large. We only displayed the peak cluster in the amygdala (see also Table 5). (B) Relationship between the connectivity and rejection rates. The mean rejection rate has no significant correlation with the connectivity (robust linear regression, slope = −0.057, *p* = 0.509 (n.s.)). (C) the connectivity and mean response time for rejection in the DI condition. The response time for acceptance increased significantly when connectivity became stronger (robust linear regression, slope = −0.592 , *p* = 0.0230 ). (D) Conceptual diagram of our findings.

**Table 5.**
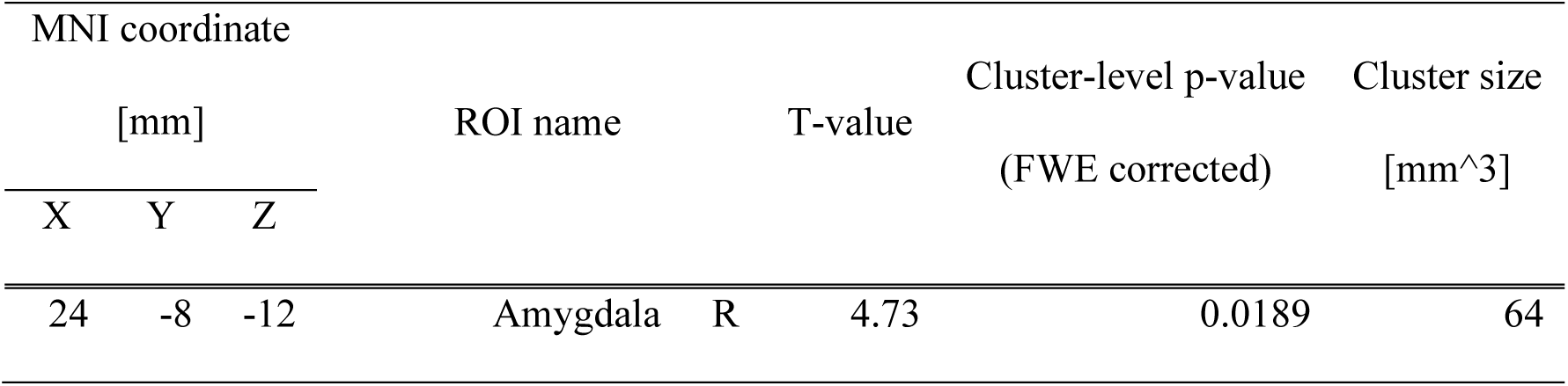
Neural activity identified by the PPI analysis whose seed ROI is the vlPFC. We conducted small volume correction for multiple comparisons (cluster-level FWE corrected *p* < 0.05) as we hypothesized bilateral amygdala is involved.

We next investigated if the interaction between the vlPFC and the amygdala encodes the rejection rate and the response times for accepting disadvantageous offers. We found that the vlPFC-amygdala connectivity exhibits no significant correlation with the rejection rate (**Fig 4B**; robust linear regression, *p* = 0.509 ) but a significant correlation with the response time for accepting DI offers (**Fig 4C**; robust linear regression, slope = −0.592 , *p* = 0.0230 ). These results suggest that the functional connectivity between the vlPFC and the amygdala is distinct from the dACC-vlPFC interaction, and that suppression from dACC to vlPFC is likely to affect the vlPFC - amygdala coupling at the offer timing.

A similar mechanism to DI may also work for SR. Therefore, we examined whether the brain activity correlated with SR at the individual level and beta values of the group correlated with −*β*(SR). We did not find significant activity (we only found activity in the dACC without multiple-comparison correction: p < 0.001, uncorrected: **S1 Fig and S1 Table**). Furthermore, this region did not show significant connectivity with any other region (uncorrected p < 0.001: only two clusters survived with a very low threshold: uncorrected p < 0.01 with a cluster-forming voxel threshold > 40 mm^3^, **S2 Table**). These observations suggest a possibility that DI has a unique and pivotal role in human bargaining behavior.

In addition to DI and SR, we also tested AI by applying the first-level results correlated with AI weighted by *β*(AI) in the second-level analysis, and we found no significant correlation (p<0.05 cluster-level FWE corrected), potentially reflecting a smaller number of AI trials.

### Comparing DDM and standard value-based models

Finally, we compared the DDM and a standard value-based model (i.e., a logistic regression model) that considers choices but not response times. We analyzed the participants’ behavior and neural activity by the logistic regression model [43]. For this purpose, we assumed that the best predictive logistic regression model has the same three parametric components as the best DDM model.

We estimated the coefficients γ of the logistic regression model using R [44] and found a significant correlation between *γ*(DI) and the mean rejection rate (Pearson’s *r* = 0.740 and *p* = 0.00328). However, the correlation between *γ*(DI) and mean response time was not significant (Pearson’s *r* = −0.0971 and *p* = 0.445). Furthermore, *γ*(DI) did not show a positive correlation with the difference in mean response times for acceptance and rejection (Pearson’s *r* = 0.0442 and *p* = 0.731, Pearson’s *r* = −0.132 and *p* = 0.345, respectively), which is in contrast to *β*(DI) (Fig. 2E **and F**). These data confirmed that only the DDM can capture behaviors in the time domain.

Observing that only *β*(DI), but not *γ*(DI), encodes response times, we compared the two parameters in the fMRI analysis. More concretely, we performed a GLM analysis, where all second-level regressors were replaced by the coefficients of the logistic model. In sharp contrast with **Fig 3A**, we identified much weaker activity in the dACC (Fig 5 **and Table 6**; p < 0.001 uncorrected). These results suggest that the DDM is more suitable for detecting the functions of the dACC in determining the choices and response times.

**Figure 5.**
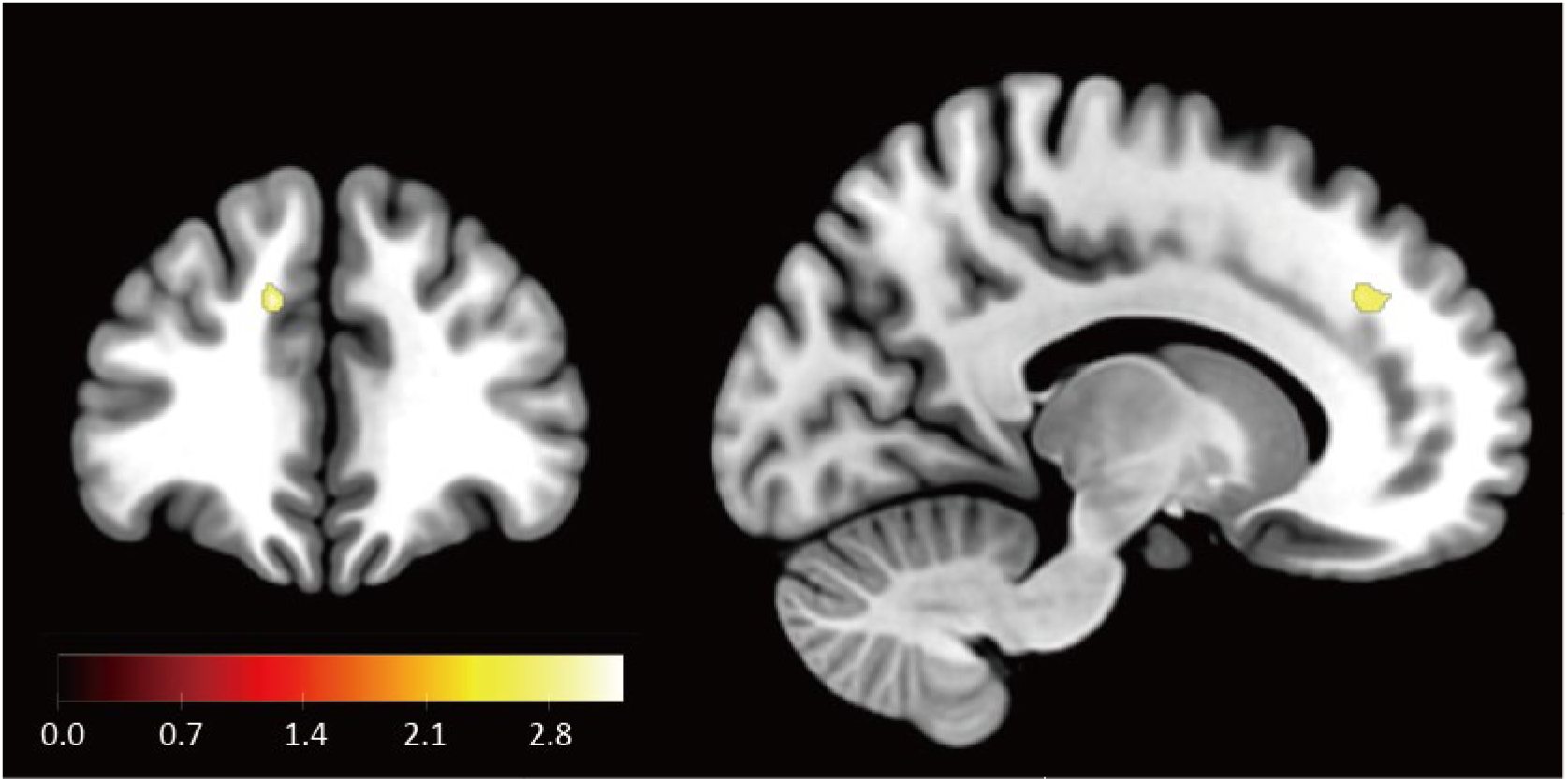
Neural activity correlated with the coefficient of the standard value-based model. We found weak neural activity in the ACC (MNI coordinate = [-12, 36, 32]; uncorrected p < 0.001; see also Table 6) and displayed the peak cluster only in the dACC (we used p < 0.005 uncorrected for display purpose).

**Table 6.**
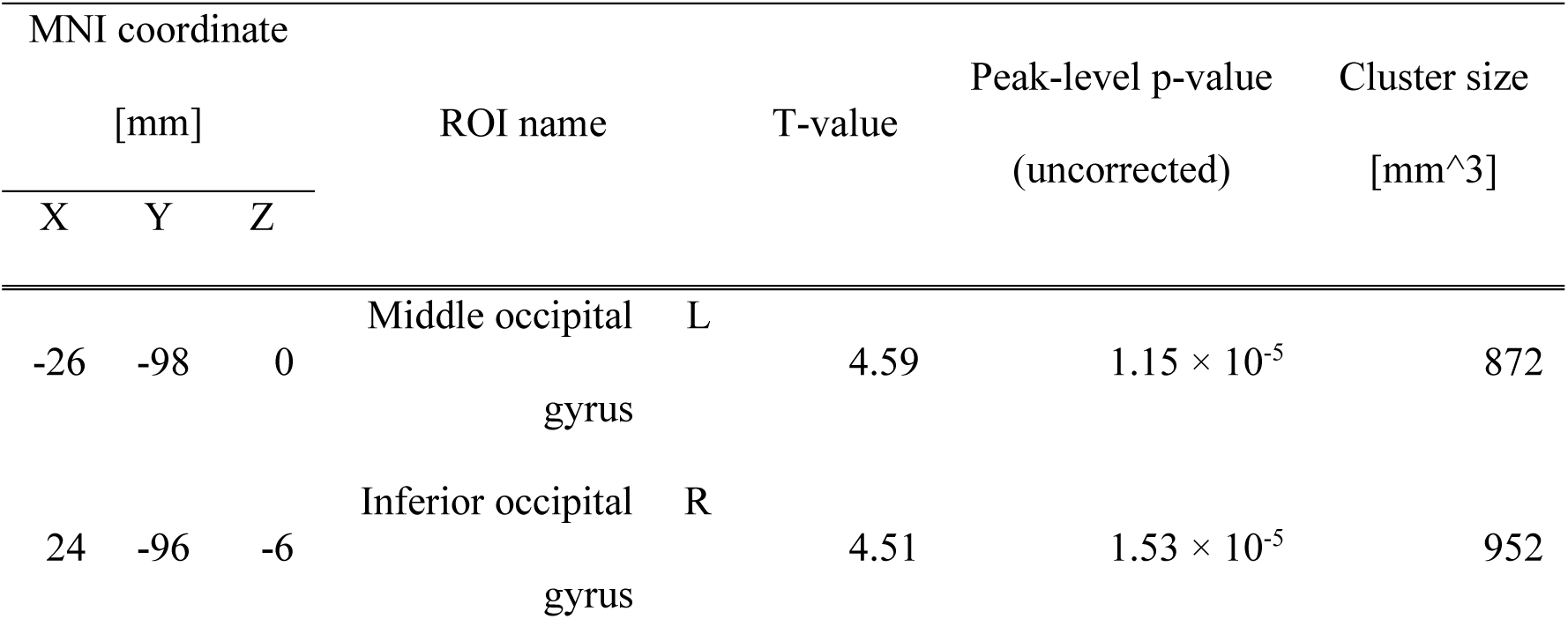
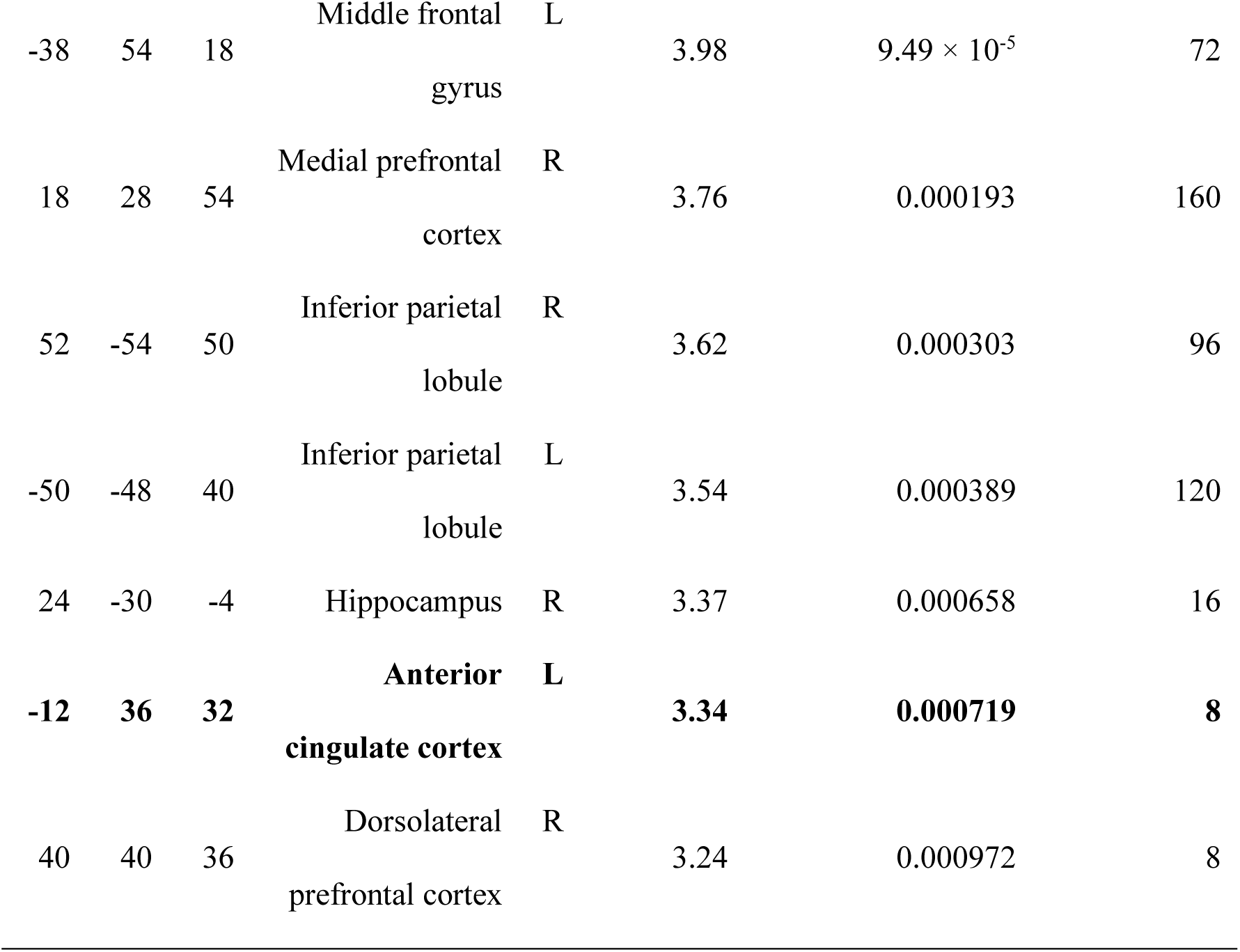
Neural activities to DI weighted by−*γ*(**DI**) **(the value-based model).** We set the statistical threshold for significance at *p* < 0.001, uncorrected for multiple comparisons and all clusters are displayed. We highlighted the neural activity in the dACC area (Peak MNI coordinate = [-12, 36, 32]).

## Discussion

The main purpose of this study was to identify the dynamic neural processes involved in accepting unfair offers. We conducted an fMRI experiment of the ultimatum game and analyzed behavioral and fMRI data using a DDM that infers internal representations to model both behavioral choices and response times. We found that participants who exhibited a smaller DDM parameter for disadvantageous inequity (DI) (i.e. *β*(DI)) and dampened disadvantage-driven rejection (i.e., higher acceptance rate) showed heightened dACC activity for DI. Our PPI analysis revealed that, when unfair offers are presented, the dACC exhibits a negative functional connectivity with the vlPFC. Importantly, we demonstrated that this PPI connectivity between dACC and vlPFC encodes the average rejection rate and the response time for accepting unfair offers. We also found that the vlPFC has a synchronized activity with the amygdala at offer timings and this connectivity is only correlated with the average response time for acceptance of unfair offers. In addition, a conventional value-based system (i.e., logistic regression) that only considers choices performed poorly compared with the DDM in detecting these fMRI signals, revealing that the consideration of both the choice and response time is key to the elucidation of the neural mechanism for accepting unfair offers.

Accepting unfair offers has been regarded as a reward-maximization process [13,45–47] since behavioral study based on choice alone favors the simplest model which can succinctly explain observed behaviors and it well suit the definition of “Economic man” [13,14,46,47]. However, the results of the present study revealed that a more complex suppression mechanism is working in the human brain to choose a reward-maximizing option without being influenced by emotion. Notably, not only the integration of response times and choices in the framework of DDM but also fMRI analysis based on the DDM parameters were central to the present findings.

We found that the PPI connectivity between the dACC and the vlPFC encodes both rejection rate and average response time for acceptance (**Fig 3E**), and that the PPI connectivity between the vlPFC and the amygdala (**Fig 4A**) encodes the average response time for acceptance, but not rejection rate. These results are consistent with previous reports that the dACC plays a pivotal role in cognitive control [31,48,49] and emotion regulation [35,41,50,51] and also that anatomically, the dACC has projections to the vlPFC [40,41,52] and the vlPFC has projections to the subcortical area, particularly the amygdala [35,40,41,51,53]. These results suggest that the dACC and the vlPFC play a central role in deciding to accept, while the link between the vlPFC and the amygdala reduce negative emotions about DI. It has been reported that the amygdala encodes economic inequity [7–9] and is involved in the rejection of unfair offers [8,10–12], suggesting that the dACC and the vlPFC suppress the negative response to inequity in the amygdala. Although our finding that activity of the amygdala and vlPFC synchronized when DI was large is consistent with this hypothesis, further investigations would be necessary to validate the hypothesis more rigorously.

This study focused on the effect of DI through dACC to achieve the acceptance of unfair (DI) offers by regulating the negative emotion caused by DI. It could be also possible that a similar suppression mechanism of SR is involved in rejecting unfair (DI and AI) offers. To address this, we conducted fMRI analysis where we looked for self-reward (SR)-correlated activity weighted by - *β*(SR). The intuition here is that when SR is large, some suppression mechanism may work to achieve rejection in those who did not take account of SR into their selection of rejection. However, we observed neither significant activity nor correlation between rejection behavior and (non-significant) weak brain activity (**S1 Fig**). These results suggest that DI has a unique impact on human bargaining behavior and that a specific suppression mechanism has developed to overcome the impact of the negative emotion for making the decision. In addition to DI and SR, advantageous inequity (AI) has also an important role. However, we did not observe a significant fMRI result. This might be because the number of trials in the AI condition are too small to detect brain activity (only 14 trials in the task).

Finally, results of the present study may also illuminate the link between bargaining behavior and well-being (including psychiatric disorders). For instance, previous research reveals that people with depression show a greater emotional response to unfair offers [54] and have a greater aversion to inequity [12,55] Furthermore, people with major depression show a lower acceptance rate for unfair offers [56] and, in healthy volunteers, inducing sadness also causes a lower acceptance rate for such offers [57]. In this context, the results of the present study may therefore contribute to deeper understanding of human well-being. Our results suggest that depression may be linked to a failure to control emotional responses to unfair offers and other attacks on pride and self-esteem. Such failure will interfere with reward-maximization.

## Materials and Methods

### Participants

We initially enlisted 71 participants for the experiment but excluded 8 due to significant or erratic head movements (greater than 1 mm) in their fMRI scans. As a result, 63 participants (41 males, 22 females; average age = 22.0, standard deviation = 3.27) were included in the main experiment. The study was approved by the ethics committee of the National Institute of Information and Communications Technology. All participants provided written informed consent.

### Experiment procedure

Participants began by reading a document outlining the task instructions and completed a consent form. They were informed that all proposals would be made by students from local universities, and that both the proposer and the proposal would vary in each trial. They were also told that their responses would affect the monetary outcomes for both the proposer and the responder. Additionally, participants were instructed to make their decisions within ten seconds and press the corresponding button. Before starting the main task, they familiarized themselves with the time limit (< 10s) and task setup through three practice trials.

### Task

Participants played the ultimatum game (**Fig 1A**) as responders. They assessed proposals from an anonymous proposer and decided whether to accept or reject the offer by pressing a button within 10 seconds. If the participant does not press the button within 10 seconds, the error trial was excluded from the analysis. If the responder accepted the offer, the money was allocated as proposed. If the responder rejected the offer, neither the proposer nor the responder received any money.

The base offers were one of the following: ¥350–150, ¥300–200, ¥250–250, ¥200–300, ¥150–350, ¥100–400, or ¥50–450 for the responder (participant) and the proposer. In each trial, a random number between ¥−25 and ¥25 was added to each reward, creating variation in the offers displayed. This design was intended to encourage participants to think critically and prevent pattern-based responses. Each base offer had eight trials, totaling 56 trials for the entire task.

Each trial began with three seconds of fixation, followed by a 1-second display of the proposer’s face and name. Participants then had 10 seconds to decide whether to accept or reject the proposal (choice period in **Fig 1A**). After making a decision, participants saw their choice displayed for 1 second (“Feedback” phase in **Fig 1A**). After the feedback, there was a 6.5-second rest period before the next trial. Each trial lasted between 22.5 and 24.5 seconds, with the entire task taking 22 minutes and 16 seconds. During the main experiment, the viewing distance was between 91.0 and 94.0 cm, the viewing angle was 21.7-22.4 degrees, and the screen luminance was 110.3 cd/m².

### Computational models for inequity-aversion

We defined disadvantageous inequity (DI) and advantageous inequity (AI) for trial t as follows:

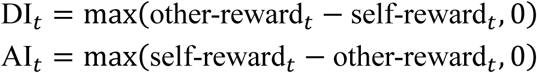

where self-reward_*t*_ ( SR_*t*_ ) represents the responder’s (participant’s) reward and other-reward_*t*_ (OR_*t*_) represents the proposer’s reward. To better understand participants’ behavior, we examined how DI, AI, and SR influenced their rejection of unfair offers and response times by applying two computational models: a drift-diffusion model (DDM) and a logistic regression model.

The DDM consists of four parameters: decision boundary, relative starting point, non-decision time, and drift rate (δ) (**Fig 2A**; For more details, see also [58]). In this study, we linked rejection to the upper boundary of the DDM and acceptance to the lower boundary. We defined δ as:

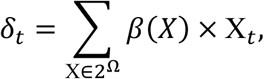

where β(“X”) represents the weight of the variable “X”, and 2^Ω refers to the power set of variables in the task, where Ω = {SR, AI, DI}. We estimated the DDM parameters using the HSSM package (https://lnccbrown.github.io/HSSM). This toolbox applies approximate hierarchical Bayesian inference to DDMs using Python and is an enhanced version of HDDM [43,44], which is commonly used for similar applications [20,59–62].

We specified a prior distribution for the DDM parameters as follows:

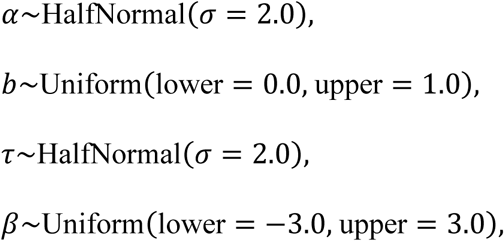

where α represents boundary separation, b represents bias, τ represents non-decision time, and β represents the linear coefficient in the drift term. Markov chain Monte Carlo (MCMC) sampling was performed with 5 chains and 500 burn-in iterations, and 500 posterior samples were obtained for each chain.

By contrast, for implementing the logistic regression model. we defined the subjective values for rejection, *V*_{*t*, reject}_, and for acceptance, *V*_{*t*, accept}_, at trial t. These values were integrated into a difference function, v(t), for the participant as follows:

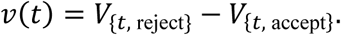

This value function was used to calculate the probability of choosing rejection, p_reject(t), using a logistic function: *p*_reject_(t) = 1/1[1 + exp(−*v*(t))].

To compare the DDM with the value-based logistic regression model, we assumed that the best predictive logistic model included the same components as the drift term in the best-performing DDM. Thus, we redefined the subjective values for rejection and acceptance as follows:

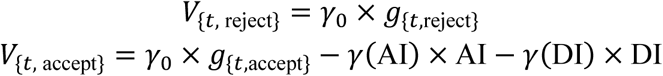

where *g*_{*t*,X}_ represents the participant’s gain for option X, and γ denotes a coefficient. In our task, *g*_{*t*,reject}_ was 0, and *g*_{*t*,accept}_ was SR_*t*_ . Logistic regression analysis was performed using the bias-reduction method provided by the Brglm package [63].

### Model selection

Since we defined the drift rate δ_*t*_ as a linear combination of the task variables, we considered nine possible models:

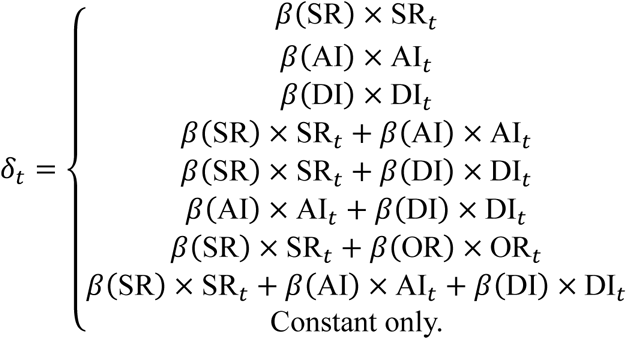

We compared the models using the widely applicable information criterion (WAIC) [37,64], a standard criterion for MCMC sampling. We computed the WAIC for each trial and summed the results to identify the model with the lowest total, which was considered the best model.

### fMRI acquisition

We assessed participants’ neural activity during task performance using fMRI. We acquired brain images with a 3T MRI (Prisma; Siemens Medical Systems) equipped with a 64-channel head coil at the Center for Information and Neural Networks, National Institute of Information and Communications Technology. To correct for distortion, we first obtained field maps covering the entire brain with the following parameters: repetition time (TR) = 753 ms, echo time 1 (TE1) = 5.16 ms, echo time 2 (TE2) = 7.62 ms, flip angle (FA) = 9°, field of view (FOV) = 256 mm, and voxel size (VS) = 2×2×2 mm. Next, we acquired T2*-weighted images during the task using a multi-echo planar pulse sequence with the parameters: TR = 2000 ms, TE = 30 ms, FA = 75°, FOV = 200 mm, VS = 2×2×2 mm, and multi-band factor is three. Finally, we obtained high-resolution T1-weighted images after the task session using the MPRAGE pulse sequence with the parameters: TR = 1900 ms, TE = 3.37 ms, FA = 9°, FOV = 256 mm, and VS = 1×1×1 mm.

### Image preprocessing

The images were processed using SPM12 (http://www.fil.ion.ucl.ac.uk/spm) within MATLAB 2018a. The preprocessing steps included slice-timing correction (with a reference time of 1000 ms), motion correction, removal of movement-induced variance (unwarping) using a measured field map, co-registration to the T1 image, spatial normalization, and spatial smoothing with a 6 mm full width at half maximum (FWHM). These procedures were carried out using the default parameters of SPM12, with the exception of the reference time for slice-timing correction and the FWHM size for spatial smoothing. After preprocessing, we identified 10 participants who had significant (> 1 mm) and/or erratic head movements, and they were excluded from the analysis.

### General linear model analysis

We conducted both individual-level and group-level general linear model (GLM) analyses. In the individual-level analysis, we created a design matrix with 14 regressors. The first regressor represented the timing of the offer event onset. The second to fourth regressors served as parametric modulators: DI, SR, and AI. The fifth regressor indicated the onset timing of the proposer’s name and face presentation. The sixth and seventh regressors represented the timing of the button press and feedback. The eighth through thirteenth regressors modeled the participant’s head movements, while the fourteenth regressor was a constant term. The event durations for the first to fourth regressors were set to the participant’s response time for each trial, while the fifth to seventh regressors had an event duration of 0 seconds. We performed the GLM analysis with these design matrices using the following default settings: a high-pass filter cutoff of 128 s, and a canonical hemodynamic response function for the basis function.

For the group-level analysis, we used contrast images related to the parametric modulator of DI from the individual-level analysis. We constructed a factorial design to explore neural activity associated with DI and analyzed the results with cluster-level family-wise error (FWE) correction at p < 0.05 (with a cluster-forming height threshold of p < 0.001).

Additionally, we created two multiple regression models to examine neural activity related to the parameters of the DDM. The first model included the participant’s β(“DI”) and β(“SR”), while the second model included γ(“DI”) and γ(“SR”). An intercept was included in each model, and the covariates were not standardized. We investigated the results using cluster-level FWE correction at p < 0.05 (with a cluster-forming height threshold of p < 0.001).

### Functional connectivity analysis

To uncover the neural mechanisms for accepting DI offers, we examined functional connectivity using seed regions derived from the peaks identified in **Fig 3A**. This analysis was conducted using the generalized psycho-physiological interaction (gPPI) toolbox [39] in SPM12. We first extracted the BOLD time series from participant-level volumes of interest (VOIs) identified in the GLM results. Each VOI was defined as a sphere with a 3 mm radius centered on the identified peak positions. The gPPI toolbox constructed PPI variables by multiplying the VOI time series with each GLM regressor (e.g., DI). Statistical significance was determined using a threshold of *p* < 0.05, corrected for multiple comparisons at the cluster level, with a cluster-forming height threshold of *p* < 0.001.

To investigate suppression from the dACC, we created a participant-level gPPI design using a seed ROI at coordinates [-10, 34, 32] (the main peak of the dACC cluster in **Fig 3A**) based on the GLM design matrix. We performed participant-level PPI analyses for DI, followed by group-level analyses using one-sample *t*-tests with a negative (i.e., - 1) contrast.

Next, we conducted the same gPPI analysis to verify whether the vlPFC region showed functional connectivity with the amygdala. For this, we constructed a participant-level PPI design using a seed ROI at coordinates [38, 44, 0] (the main peak of the vlPFC cluster in **Fig 3C**) multiplied by DI. Participant-level gPPI analyses were performed to identify regions showing synchronized activity with the vlPFC at the time of the offer. Group-level PPI analysis was conducted using one-sample *t*-tests with positive contrasts (i.e., 1). Given our a priori hypothesis of an interaction between the vlPFC and the amygdala, we created target masks for the left and right amygdala using the Harvard-Oxford subcortical structural atlas, applying an 80% probability threshold for voxel inclusion in the amygdala. Small volume correction for multiple comparisons was applied (cluster-level FWE-corrected, *p* < 0.05). In this analysis, two participants were excluded due to signal dropout in their amygdala regions on fMRI images.

### BOLD signal time-series analysis

We obtained BOLD signal time-series by calculating peri-stimulus time histogram (PSTH) at the target position. We first constructed a design matrix that consisted of 15 regressors for each. The first to seventh regressors represented the offer timing in each condition. The eighth to thirteenth regressors represented the participant’s head movement, and the fourteenth was a constant. The event duration of the first to the seventh regressor was set to a participant’s reaction time in each trial. We conducted individual GLM analysis with this design matrix and obtained each participant’s PSTH at the dACC (MNI coordinates are [-10, 34, 32]) from this GLM result. We then calculated the average of PSTHs across 5 + 5 participants who had the smallest and largest β(envy)s, respectively.

## Supporting Information

**S1 Figure.**
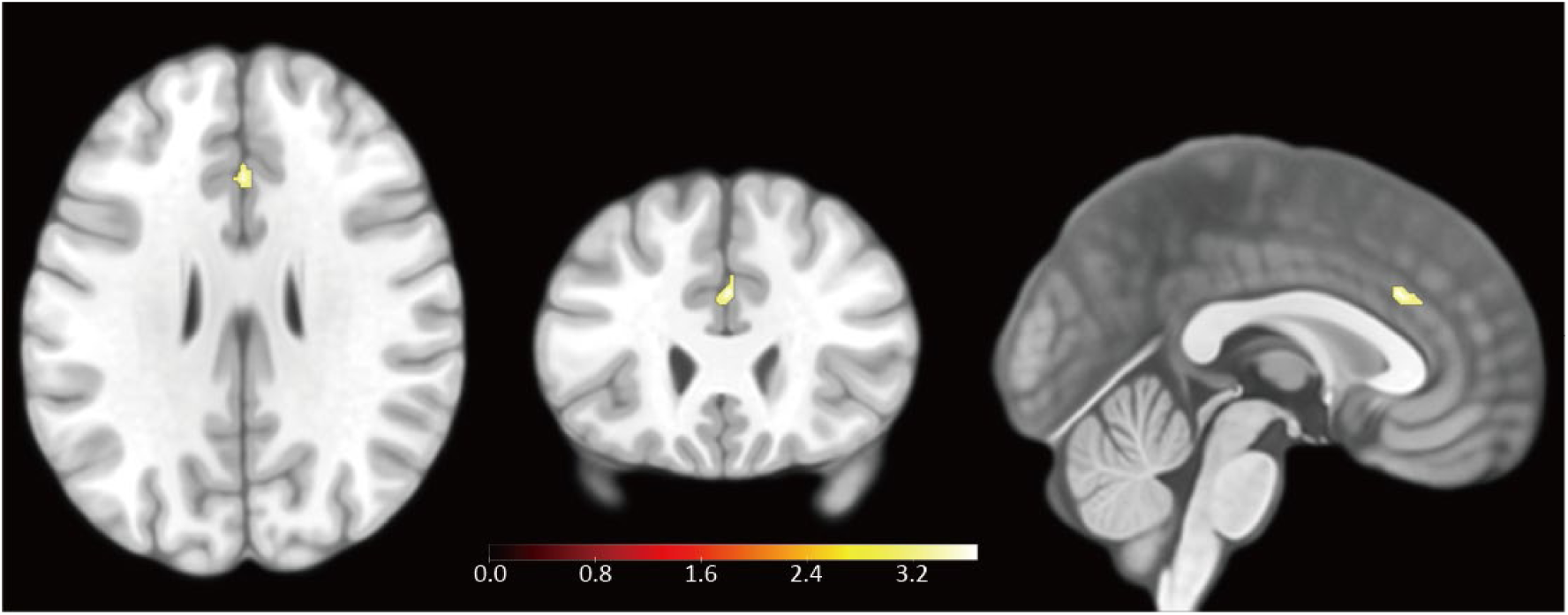
Neural activity correlated with SR weighted by - *β*(SR) We found non-significant neural activity in the ACC (Peak MNI coordinate = [0, 26, 28]; uncorrected *p* < 0.001).

**S1 Table.**
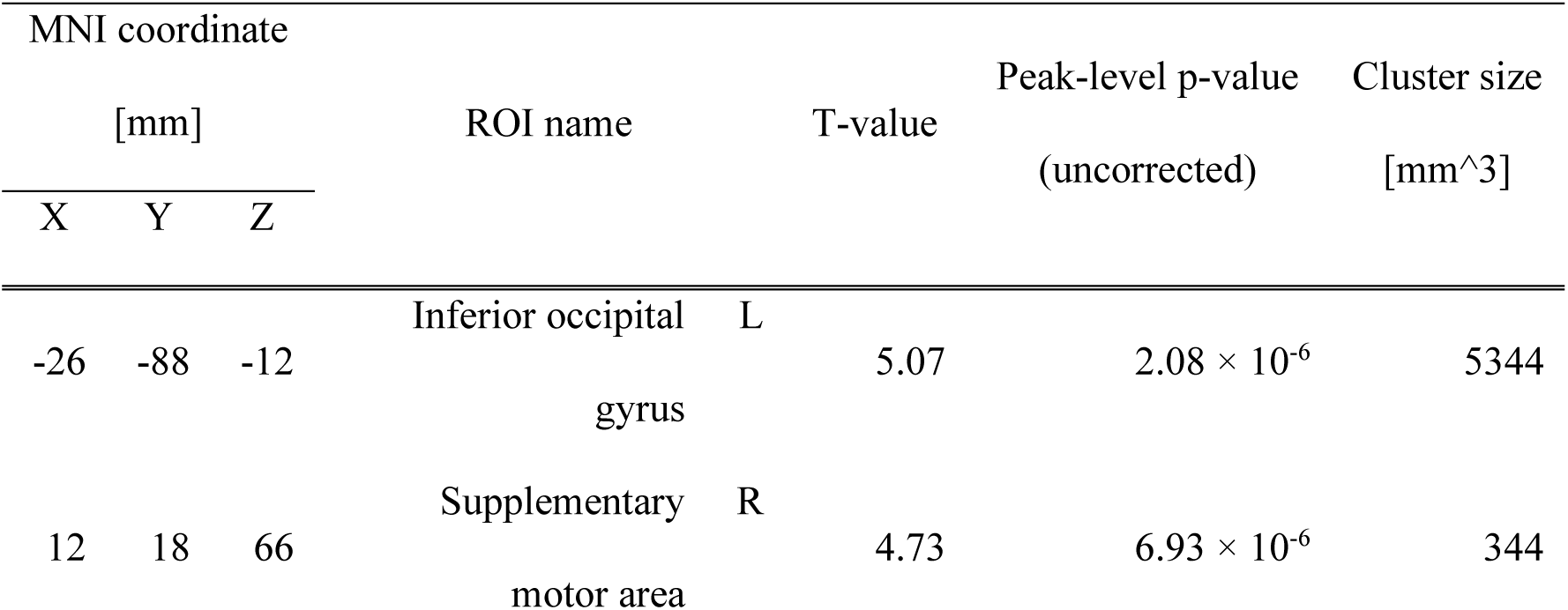
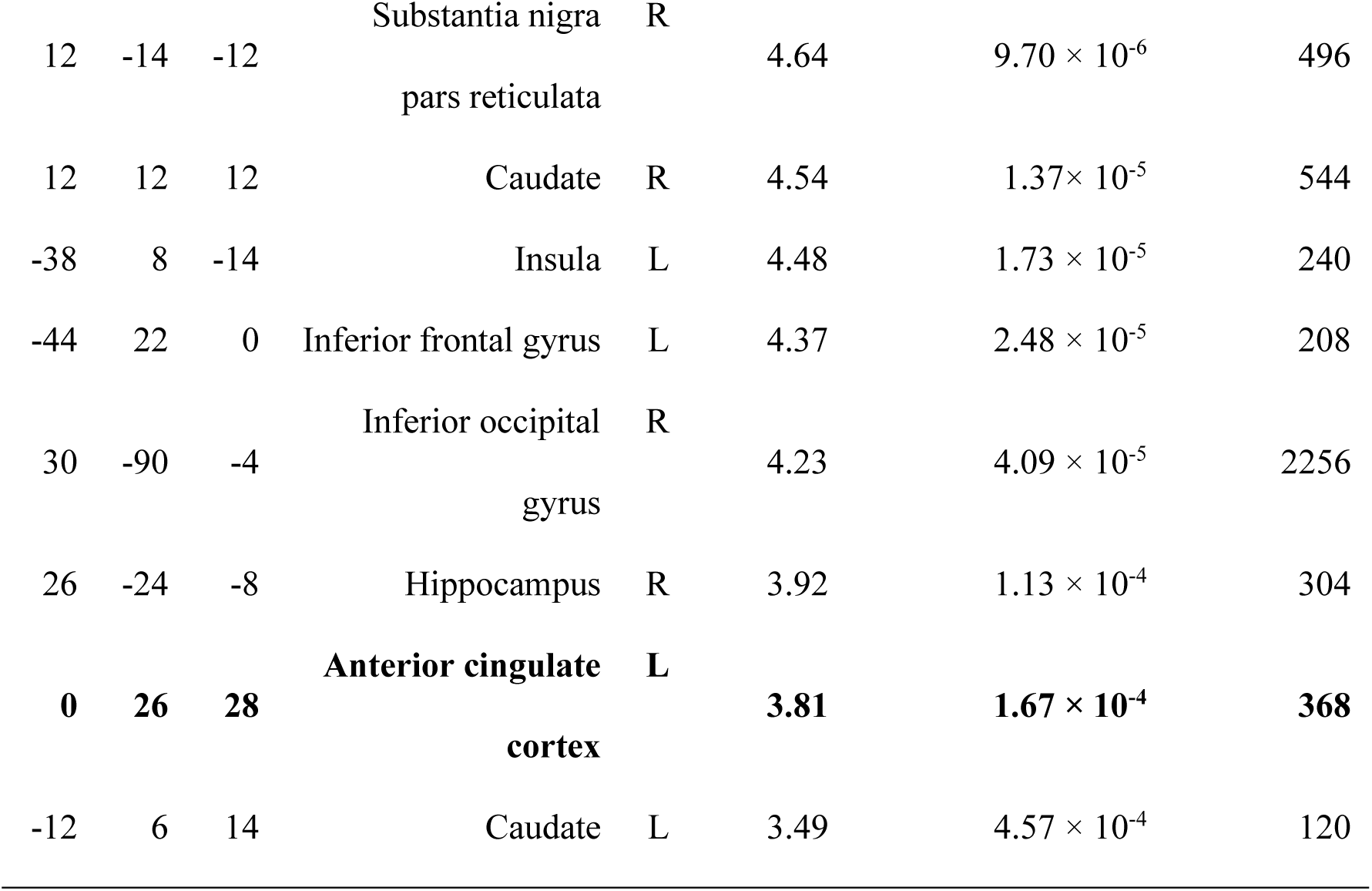
List of neural activity with SR weighted by. −*β*(**SR**). We set the statistical threshold for significance at *p* < 0.001, uncorrected for multiple comparisons and clusters larger than 80 mm^3^ are displayed.

**S2 Table.**
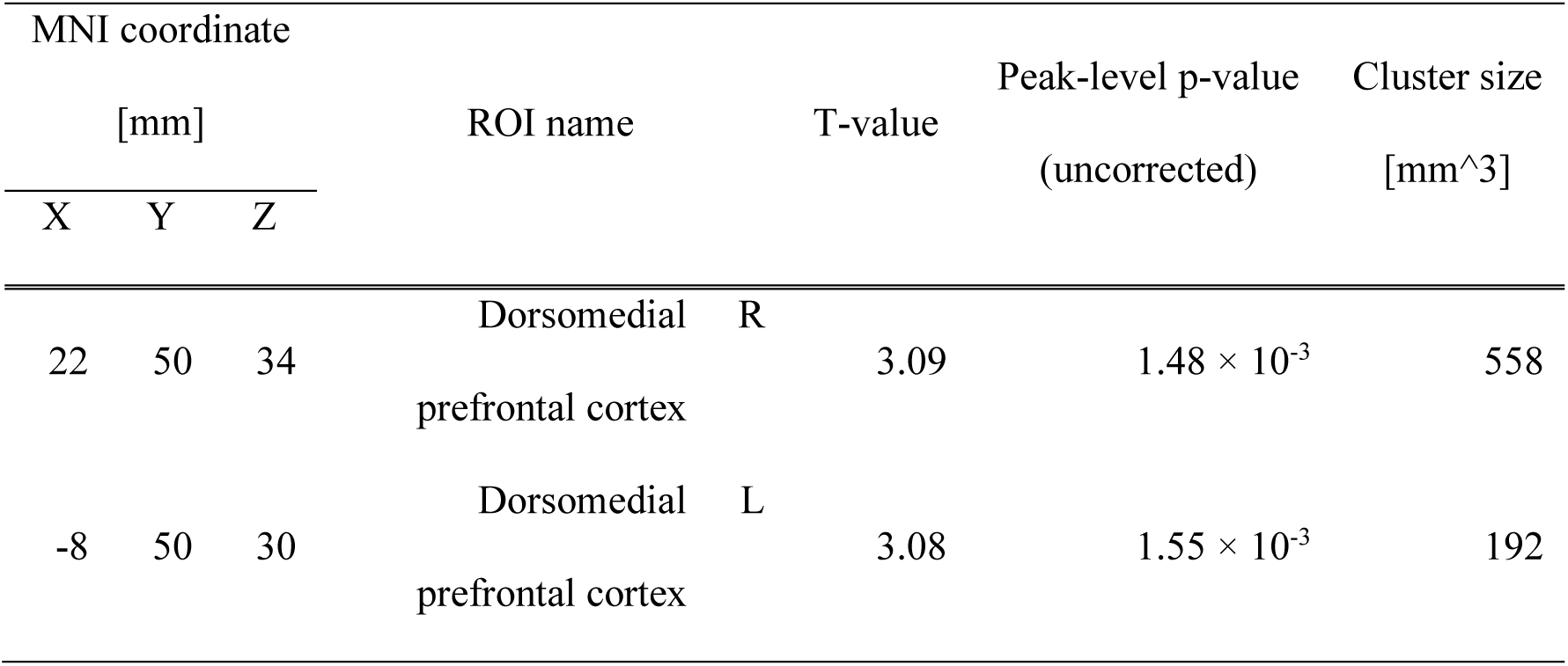
Neural activity identified by the PPI analysis whose seed ROI is the ACC in S1 Table. We set the statistical threshold for significance at *p* < 0.001, uncorrected for multiple comparisons and clusters larger than 80 mm^3^ are displayed.

## Reference

1. Fehr E, Gächter S. Fairness and Retaliation: The Economics of Reciprocity. SSRN Electron J. 2000. doi:10.2139/ssrn.229149

2. Henrich J, Boyd R, Bowles S, Camerer C, Fehr E, Gintis H, et al. In Search of Homo Economicus: Behavioral Experiments in 15 Small-Scale Societies. Am Econ Rev. 2001;91: 73–78. doi:10.1257/aer.91.2.73

3. Sanfey AG, Rilling JK, Aronson JA, Nystrom LE, Cohen JD. The Neural Basis of Economic Decision-Making in the Ultimatum Game. Science. 2003;300: 1755–1758. doi:10.1126/science.1082976

4. Gu X, Wang X, Hula A, Wang S, Xu S, Lohrenz TM, et al. Necessary, Yet Dissociable Contributions of the Insular and Ventromedial Prefrontal Cortices to Norm Adaptation: Computational and Lesion Evidence in Humans. J Neurosci. 2015;35: 467–473. doi:10.1523/jneurosci.2906-14.2015

5. Güroğlu B, Bos W van den, Rombouts SARB, Crone EA. Unfair? It depends: Neural correlates of fairness in social context. Soc Cogn Affect Neur. 2010;5: 414–423. doi:10.1093/scan/nsq013

6. Gabay AS, Radua J, Kempton MJ, Mehta MA. The Ultimatum Game and the brain: A meta-analysis of neuroimaging studies. Neurosci Biobehav Rev. 2014;47: 549–558. doi:10.1016/j.neubiorev.2014.10.014

7. Haruno M, Frith CD. Activity in the amygdala elicited by unfair divisions predicts social value orientation. Nat Neurosci. 2010;13: 160–161. doi:10.1038/nn.2468

8. Gospic K, Mohlin E, Fransson P, Petrovic P, Johannesson M, Ingvar M. Limbic Justice—Amygdala Involvement in Immediate Rejection in the Ultimatum Game. PLoS Biol. 2011;9: e1001054. doi:10.1371/journal.pbio.1001054

9. Haruno M. Unraveling how the third-party brain under stress responds to injustices. PLOS Biol. 2024;22: e3002618. doi:10.1371/journal.pbio.3002618

10. Haruno M, Kimura M, Frith CD. Activity in the Nucleus Accumbens and Amygdala Underlies Individual Differences in Prosocial and Individualistic Economic Choices. J Cognitive Neurosci. 2014;26: 1861–1870. doi:10.1162/jocn_a_00589

11. Scheele D, Mihov Y, Kendrick KM, Feinstein JS, Reich H, Maier W, et al. Amygdala Lesion Profoundly Alters Altruistic Punishment. Biol Psychiatry. 2012;72: e5–e7. doi:10.1016/j.biopsych.2012.01.028

12. Tanaka T, Yamamoto T, Haruno M. Brain response patterns to economic inequity predict present and future depression indices. Nat Hum Behav. 2017;1: 748–756. doi:10.1038/s41562-017-0207-1

13. Fehr E, Schmidt KM. A Theory of Fairness, Competition, and Cooperation. Q J Econ. 1999;114: 817–868. doi:10.1162/003355399556151

14. Güth W, Schmittberger R, Schwarze B. An experimental analysis of ultimatum bargaining. J Econ Behav Organ. 1982;3: 367–388. doi:10.1016/0167-2681(82)90011-7

15. Krajbich I, Hare T, Bartling B, Morishima Y, Fehr E. A Common Mechanism Underlying Food Choice and Social Decisions. PLoS Comput Biol. 2015;11: e1004371. doi:10.1371/journal.pcbi.1004371

16. Rand DG, Greene JD, Nowak MA. Spontaneous giving and calculated greed. Nature. 2012;489: 427. doi:10.1038/nature11467

17. Rubinstein A. Instinctive and Cognitive Reasoning: A Study of Response Times. SSRN Electron J. 2006. doi:10.2139/ssrn.889310

18. Yamagishi T, Matsumoto Y, Kiyonari T, Takagishi H, Li Y, Kanai R, et al. Response time in economic games reflects different types of decision conflict for prosocial and proself individuals. Proc Natl Acad Sci. 2017;114: 6394–6399. doi:10.1073/pnas.1608877114

19. Kahneman D. Thinking, fast and slow. Farrar, Straus and Giroux; 2011.

20. Ratcliff R, Smith PL, Brown SD, McKoon G. Diffusion Decision Model: Current Issues and History. Trends Cogn Sci. 2016;20: 260–281. doi:10.1016/j.tics.2016.01.007

21. Ratcliff R. A theory of memory retrieval. Psychol Rev. 1978;85: 59. doi:10.1037/0033-295x.85.2.59

22. Brunton BW, Botvinick MM, Brody CD. Rats and Humans Can Optimally Accumulate Evidence for Decision-Making. Science. 2013;340: 95–98. doi:10.1126/science.1233912

23. Kiani R, Shadlen MN. Representation of Confidence Associated with a Decision by Neurons in the Parietal Cortex. Science. 2009;324: 759–764. doi:10.1126/science.1169405

24. Ratcliff R, Cherian A, Segraves M. A Comparison of Macaque Behavior and Superior Colliculus Neuronal Activity to Predictions From Models of Two-Choice Decisions. J Neurophysiol. 2003;90: 1392–1407. doi:10.1152/jn.01049.2002

25. Shadlen MN, Newsome WT. Motion perception: seeing and deciding. Proc National Acad Sci. 1996;93: 628–633. doi:10.1073/pnas.93.2.628

26. Ratcliff R, Rouder JN. A Diffusion Model Account of Masking in Two-Choice Letter Identification. J Exp Psychol: Hum Percept Perform. 2000;26: 127–140. doi:10.1037/0096-1523.26.1.127

27. Smith PL, Ratcliff R, Wolfgang BJ. Attention orienting and the time course of perceptual decisions: response time distributions with masked and unmasked displays. Vis Res. 2004;44: 1297–1320. doi:10.1016/j.visres.2004.01.002

28. Smith PL, Ratcliff R. An Integrated Theory of Attention and Decision Making in Visual Signal Detection. Psychol Rev. 2009;116: 283–317. doi:10.1037/a0015156

29. Chen F, Krajbich I. Biased sequential sampling underlies the effects of time pressure and delay in social decision making. Nat Commun. 2018;9: 3557. doi:10.1038/s41467-018-05994-9

30. Hutcherson CA, Bushong B, Rangel A. A Neurocomputational Model of Altruistic Choice and Its Implications. Neuron. 2015;87: 451–462. doi:10.1016/j.neuron.2015.06.031

31. Shackman AJ, Salomons TV, Slagter HA, Fox AS, Winter JJ, Davidson RJ. The integration of negative affect, pain and cognitive control in the cingulate cortex. Nat Rev Neurosci. 2011;12: 154–167. doi:10.1038/nrn2994

32. Pruessner L, Barnow S, Holt DV, Joormann J, Schulze K. A Cognitive Control Framework for Understanding Emotion Regulation Flexibility. Emotion. 2020;20: 21–29. doi:10.1037/emo0000658

33. Inzlicht M, Bartholow BD, Hirsh JB. Emotional foundations of cognitive control. Trends Cogn Sci. 2015;19: 126–132. doi:10.1016/j.tics.2015.01.004

34. Ochsner KN, Silvers JA, Buhle JT. Functional imaging studies of emotion regulation: a synthetic review and evolving model of the cognitive control of emotion. Ann N York Acad Sci. 2012;1251: E1–E24. doi:10.1111/j.1749-6632.2012.06751.x

35. Ochsner KN, Gross JJ. The cognitive control of emotion. Trends Cogn Sci. 2005;9: 242–249. doi:10.1016/j.tics.2005.03.010

36. Loewenstein GF, Thompson L, Bazerman MH. Social Utility and Decision Making in Interpersonal Contexts. J Pers Soc Psychol. 1989;57: 426–441. doi:10.1037/0022-3514.57.3.426

37. Watanabe S. Asymptotic Equivalence of Bayes Cross Validation and Widely Applicable Information Criterion in Singular Learning Theory. Journal of Machine Learning Research. 2010;11: 3571--3594. Available: http://jmlr.org/papers/v11/watanabe10a.html

38. Gelman A, Carlin JB, Stern HS, Dunson DB, Vehtari A, Rubin DB. Bayesian Data Analysis. 2019. doi:10.1201/b16018

39. McLaren DG, Ries ML, Xu G, Johnson SC. A generalized form of context-dependent psychophysiological interactions (gPPI): A comparison to standard approaches. Neuroimage. 2012;61: 1277–1286. doi:10.1016/j.neuroimage.2012.03.068

40. Berboth S, Morawetz C. Amygdala-prefrontal connectivity during emotion regulation: A meta-analysis of psychophysiological interactions. Neuropsychologia. 2021;153: 107767. doi:10.1016/j.neuropsychologia.2021.107767

41. Haber SN, Liu H, Seidlitz J, Bullmore E. Prefrontal connectomics: from anatomy to human imaging. Neuropsychopharmacol. 2022;47: 20–40. doi:10.1038/s41386-021-01156-6

42. Buhle JT, Silvers JA, Wager TD, Lopez R, Onyemekwu C, Kober H, et al. Cognitive Reappraisal of Emotion: A Meta-Analysis of Human Neuroimaging Studies. Cereb Cortex. 2014;24: 2981–2990. doi:10.1093/cercor/bht154

43. Bellemare C, Kröger S, Soest AV. Measuring Inequity Aversion in a Heterogeneous Population Using Experimental Decisions and Subjective Probabilities. Econometrica. 2008;76: 815–839. doi:10.1111/j.1468-0262.2008.00860.x

44. Team RC. R: A Language and Environment for Statistical Computing. 2023. Available: https://www.R-project.org/

45. Tabibnia G, Satpute AB, Lieberman MD. The Sunny Side of Fairness. Psychol Sci. 2007;19: 339–347. doi:10.1111/j.1467-9280.2008.02091.x

46. Bolton GE, Zwick R. Anonymity versus Punishment in Ultimatum Bargaining. Games Econ Behav. 1995;10: 95–121. doi:10.1006/game.1995.1026

47. Henrich J, Boyd R, Bowles S, Camerer C, Fehr E, Gintis H, et al. “Economic man” in cross-cultural perspective: Behavioral experiments in 15 small-scale societies. Behav Brain Sci. 2005;28: 795–815. doi:10.1017/s0140525x05000142

48. Shenhav A, Botvinick MM, Cohen JD. The Expected Value of Control: An Integrative Theory of Anterior Cingulate Cortex Function. Neuron. 2013;79: 217–240. doi:10.1016/j.neuron.2013.07.007

49. Petersen SE, Posner MI. The Attention System of the Human Brain: 20 Years After. Annu Rev Neurosci. 2012;35: 73–89. doi:10.1146/annurev-neuro-062111-150525

50. Langner R, Leiberg S, Hoffstaedter F, Eickhoff SB. Towards a human self-regulation system: Common and distinct neural signatures of emotional and behavioural control. Neurosci Biobehav Rev. 2018;90: 400–410. doi:10.1016/j.neubiorev.2018.04.022

51. Etkin A, Egner T, Kalisch R. Emotional processing in anterior cingulate and medial prefrontal cortex. Trends Cogn Sci. 2011;15: 85–93. doi:10.1016/j.tics.2010.11.004

52. Monosov IE, Rushworth MFS. Interactions between ventrolateral prefrontal and anterior cingulate cortex during learning and behavioural change. Neuropsychopharmacology. 2022;47: 196–210. doi:10.1038/s41386-021-01079-2

53. Sakagami M, Pan X. Functional role of the ventrolateral prefrontal cortex in decision making. Curr Opin Neurobiol. 2007;17: 228–233. doi:10.1016/j.conb.2007.02.008

54. Nicolaisen-Sobesky E, Paz V, Cervantes-Constantino F, Fernández-Theoduloz G, Pérez A, Martínez-Montes E, et al. Event-related potentials during the ultimatum game in people with symptoms of depression and/or social anxiety. Psychophysiology. 2023;60: e14319. doi:10.1111/psyp.14319

55. Pulcu E, Thomas EJ, Trotter PD, McFarquhar M, Juhasz G, Sahakian BJ, et al. Social-economical decision making in current and remitted major depression. Psychol Med. 2015;45: 1301–1313. doi:10.1017/s0033291714002414

56. Jin Y, Gao Q, Wang Y, Xiao L, Wu MS, Zhou Y. The Perception-Behavior Dissociation in the Ultimatum Game in Unmedicated Patients With Major Depressive Disorders. J Psychopathol Clin Sci. 2022;131: 253–264. doi:10.1037/abn0000747

57. Harlé KM, Sanfey AG. Incidental Sadness Biases Social Economic Decisions in the Ultimatum Game. Emotion. 2007;7: 876–881. doi:10.1037/1528-3542.7.4.876

58. Numano S, Haruno M. Detailed analysis of drift diffusion model parameters estimated for the ultimatum game. Neurosci Res. 2024. doi:10.1016/j.neures.2024.12.003

59. Ratcliff R, Childers R. Individual Differences and Fitting Methods for the Two-Choice Diffusion Model of Decision Making. Decis. 2015;2: 237–279. doi:10.1037/dec0000030

60. Westbrook A, Bosch R van den, Määttä JI, Hofmans L, Papadopetraki D, Cools R, et al. Dopamine promotes cognitive effort by biasing the benefits versus costs of cognitive work. Science. 2020;367: 1362–1366. doi:10.1126/science.aaz5891

61. Rollwage M, Loosen A, Hauser TU, Moran R, Dolan RJ, Fleming SM. Confidence drives a neural confirmation bias. Nat Commun. 2020;11: 2634. doi:10.1038/s41467-020-16278-6

62. Pagnier GJ, Asaad WF, Frank MJ. Double dissociation of dopamine and subthalamic nucleus stimulation on effortful cost/benefit decision making. Curr Biol. 2024;34: 655–660.e3. doi:10.1016/j.cub.2023.12.045

63. Kosmidis I, Firth D. Jeffreys-prior penalty, finiteness and shrinkage in binomial-response generalized linear models. Biometrika. 2020;108: 71–82. doi:10.1093/biomet/asaa052

64. Spiegelhalter DJ, Best NG, Carlin BP, Linde AVD. Bayesian measures of model complexity and fit. J R Stat Soc: Ser B (Stat Methodol). 2002;64: 583–639. doi:10.1111/1467-9868.00353

